# The Bacterial Quorum-Sensing Signal 2-Aminoacetophenone Rewires Immune Cell Bioenergetics through the PGC-1α/ERRα Axis to Mediate Tolerance to Infection

**DOI:** 10.1101/2024.02.26.582124

**Authors:** Arijit Chakraborty, Arunava Bandyopadhaya, Vijay K Singh, Filip Kovacic, Sujin Cha, William M. Oldham, A. Aria Tzika, Laurence G Rahme

**Affiliations:** Department of Surgery, Massachusetts General Hospital, and Harvard Medical School, Boston, Massachusetts, USA; Shriners Hospitals for Children Boston, Boston, Massachusetts, USA; Department of Microbiology, Harvard Medical School, Boston, Massachusetts, USA; Institute of Molecular Enzyme Technology, Heinrich Heine University Düsseldorf, Forschungszentrum Jülich GmbH, Jülich, Germany; Department of Medicine, Brigham and Women’s Hospital and Harvard Medical School, Boston, Massachusetts, USA

**Author notes:** **Corresponding author:** Laurence G. Rahme, Ph.D (LGR), Address: 340 Their Research Building, 50 Blossom Street, Massachusetts General Hospital, Boston, MA 02114. These authors contributed equally (Arijit Chakraborty, Arunava Bandyopadhaya).

**Keywords:** Pseudomonas aeruginosa, quorum sensing, infection, macrophages, bacterial persistence, 2-aminoacetophenone (2-AA), MvfR, PqsR, immunometabolism, host tolerization, histone deacetylation, epigenetic reprogramming, metabolic reprogramming, oxidative phosphorylation, pyruvate, mitochondrial pyruvate carrier (MPC1), peroxisome proliferator-activated receptor gamma co-activator 1-alpha (PGC-1α), estrogen-related receptor (ERRα), acetyl-CoA, ATP, bioenergetics

## Abstract

How bacterial pathogens exploit host metabolism to promote immune tolerance and persist in infected hosts remains elusive. To achieve this, we show that *Pseudomonas aeruginosa (PA),* a recalcitrant pathogen, utilizes the quorum sensing (QS) signal 2-aminoacetophenone (2-AA). Here, we unveil how 2-AA-driven immune tolerization causes distinct metabolic perturbations in macrophages’ mitochondrial respiration and bioenergetics. We present evidence indicating that these effects stem from decreased pyruvate transport into mitochondria. This reduction is attributed to decreased expression of the mitochondrial pyruvate carrier (MPC1), which is mediated by diminished expression and nuclear presence of its transcriptional regulator, estrogen-related nuclear receptor alpha (ERRα). Consequently, ERRα exhibits weakened binding to the MPC1 promoter. This outcome arises from the impaired interaction between ERRα and the peroxisome proliferator-activated receptor gamma coactivator 1-alpha (PGC-1α). Ultimately, this cascade results in diminished pyruvate influx into mitochondria and, consequently reduced ATP production in tolerized macrophages. Exogenously added ATP in infected macrophages restores the transcript levels of *MPC1* and *ERR-*α *and* enhances cytokine production and intracellular bacterial clearance. Consistent with the *in vitro* findings, murine infection studies corroborate the 2-AA-mediated long-lasting decrease in ATP and acetyl-CoA and its association with *PA* persistence, further supporting this QS signaling molecule as the culprit of the host bioenergetic alterations and *PA* persistence. These findings unveil 2-AA as a modulator of cellular immunometabolism and reveal an unprecedented mechanism of host tolerance to infection involving the PGC-1α/ERRα axis in its influence on MPC1/OXPHOS-dependent energy production and *PA* clearance. These paradigmatic findings pave the way for developing treatments to bolster host resilience to pathogen-induced damage. Given that QS is a common characteristic of prokaryotes, it is likely that 2-AA-like molecules with similar functions may be present in other pathogens.

## Introduction

Host tolerance is a fundamental mechanism of innate immunity. Studies have shown that innate immune cells following infection or exposure to microbial products may enter a state of immune tolerance characterized by reduced responsiveness towards microbial re-exposure a few hours later [1–3]. Epigenetic and signaling-based mechanisms have been proposed to be involved in host immune tolerance [1–3]. In recent years, several reports have highlighted the complex interplay between metabolic reprogramming and immunity [4, 5]. It was suggested that specific metabolic programs are activated in monocytes and macrophages upon exposure to infection and microbial products [6, 7]. However, in host tolerance, the molecular mechanisms underlying immunometabolic reprogramming mediated by bacterial pathogens remain poorly understood.

Metabolic pathways are essential for generating energy for various cellular functions, including those performed by immune cells [8, 9]. Macrophages use metabolic pathways to generate energy and metabolites to adapt to changing environments and stimuli, thereby enabling them to cope with the needs of a fluctuating immune response [7, 10]. Thus, properly functioning metabolic pathways are vital for immune cells to counteract pathogens. For instance, the crucial energy-carrying molecule adenosine triphosphate (ATP) is required to phagocytose pathogen-derived molecules efficiently.

*Pseudomonas aeruginosa (PA)*, a recalcitrant ESKAPE (*<u>E</u>nterococcus faecium, <u>S</u>taphylococcus aureus, <u>K</u>lebsiella pneumoniae, <u>A</u>cinetobacter baumannii, <u>P</u>A,* and *<u>E</u>nterobacter sp.*) pathogen that causes acute and persistent infections, secretes virulence-associated low molecular weight signaling molecules, several of which are regulated by quorum sensing (QS) [11–14] and able to modulate host immune responses [15–18]. QS is a cell density-dependent signaling system bacteria use to synchronize their activities [11–14]. MvfR (a.k.a. PqsR), a critical QS transcription factor of *PA,* regulates the synthesis of many small molecules, including signaling molecules such as 2-aminoacetophenone (2-AA) [11, 19–21]. *In vivo* studies have demonstrated that 2-AA enables *PA* to persist in infected murine tissues by promoting innate immune tolerance through histone deacetylase 1 (HDAC1)-mediated epigenetic reprogramming [17, 18]. 2-AA tolerization reprograms the host inflammatory signaling cascade by maintaining chromatin in a “silent” state through increased HDAC1 expression and activity and decreased histone acetyltransferase (HAT) activity [18]. These changes also impact the protein-protein interaction between HAT and cyclic AMP response element-binding protein/HDAC1 and p50/p65 nuclear factor (NF)-κB subunits [18, 22]. The 2-AA-mediated tolerization permits *PA* to persist in infected tissues [17, 18], underscoring the difference from lipopolysaccharide (LPS)-mediated tolerization, which instead leads to bacterial clearance and involves different HDACs [23].

Mitochondria, the “powerhouse” of the cells, are crucial for the regulation, differentiation, and survival of macrophages and other immune cells [8]. Our group’s previous *in vivo* and *in vitro* studies indicated that 2-AA affects metabolic functions in skeletal muscle, which contains a high concentration of mitochondria [24–26]. Injection of 2-AA in murine skeletal muscle dampens the expression of genes associated with OXPHOS and the master regulator of mitochondrial biogenesis peroxisome proliferator-activated receptor-γ coactivator-1 beta (PGC-1β) [24]. Another *PA* QS molecule, 3-oxo-C12-HSL, attenuates the expression of PGC-1α [27]. PGC-1α and PGC-1β belong to the PPARγ family of inducible transcriptional coactivators and are known regulators of mitochondrial metabolism [28]. The coactivator PGC-1 proteins are involved in various cellular energy metabolic processes [29, 30], including but not limited to mitochondrial metabolism [31]. Recent studies have also emphasized the role of estrogen-related nuclear receptor α (ERRα) in coordinating metabolic capacity with energy demand in health and disease [32]. PGC-1α activates ERRα transcriptional activity through protein-protein interaction, which enhances the expression of ERRα [33, 34] and other mitochondrial genes involved in lipid metabolism and OXPHOS [35]. PGC-1α/ERRα axis has been extensively studied in cancer and established as a central regulatory node of energy metabolism that induces the global expression of genes involved in mitochondrial biogenesis and functions [36–38]. ERRα has been shown to occupy the promoter regions of many genes participating in the TCA cycle and OXPHOS [39, 40], including the mitochondrial pyruvate carrier (MPC1). In human renal carcinoma cells, PGC-1α/ERRα interaction results in efficient activation of MPC1 expression and transport of pyruvate into mitochondrion for efficient OXPHOS and energy production [41].

Given that our previous studies pointed to the 2-AA effect on mitochondrial functions [24–26] and that mitochondria are crucial for the regulation, differentiation, and survival of macrophages and other immune cells [8], we investigated how 2-AA may mediate cellular metabolic reprogramming in tolerized immune cells by examining the link between the PGC-1α/ERRα axis and 2-AA-mediated immune tolerance. Our findings uncovered the crucial action of 2-AA on energy homeostasis and metabolism in tolerized immune cells through perturbances of the ERRα/PGC-1α interaction, MPC1-mediated pyruvate transport and the production of the key energy metabolism molecules, ATP and acetyl-CoA and their association with the persistence of *PA* in mammalian tissues.

## Results

### Tolerization by 2-AA impacts the generation of crucial energy metabolites in macrophages

Following infection or exposure to microbial products, innate immune cells may enter a state of tolerance several hours later, as depicted by their diminished responsiveness or unresponsiveness to microbial re-exposure [1–3, 42]. Given that 2-AA tolerized cells are non-responsive to 2-AA re-exposure, we investigated the impact of 2-AA on energy homeostasis and metabolism in non-tolerized and tolerized BMDM cells (Fig. 1). We quantified the levels of ATP and acetyl-CoA, which is a fuel for ATP production, in a series of *in vitro* experiments in which murine BMDM cells were exposed to 2-AA (1^st^ 2-AA exposure) or re-exposed (2^nd^ 2-AA exposure) to respectively determine their activation, and tolerance as well as memory to this QS molecule (Fig. 1A). Initially, following the 1^st^ 2-AA exposure (black bars), a significant increase in the concentrations of intracellular ATP (Fig. 1B) and acetyl-CoA (Fig. 1C) was observed in BMDM cells exposed to 2-AA for 1 and 6 h compared to their corresponding naïve control cells (grey bars). However, cells exposed to 2-AA for 48 h (black bar) appeared to have entered a state of tolerance as ATP and acetyl-CoA levels were decreased compared to 2-AA exposures for a shorter time (Figs. 1B and 1C). The tolerance of these cells as well as their memory to 2-AA was determined by their responsiveness to a 2^nd^ 2-AA exposure (Fig. 1A). To achieve this, these cells were washed to remove 2-AA and allowed to rest for 24 or 106 h in the absence of 2-AA before receiving the 2^nd^ 2-AA exposure for 1 or 6 h (Fig. 1A). Figures 1B and 1C (red bars) show the unresponsiveness of BMDM cells to 2-AA re-exposure as evidenced by the similar levels of ATP and acetyl-CoA compared to cells exposed to 2-AA for 48 h (1^st^ exposure black bars). The sustained unresponsiveness of the cells to re-exposure indicates that the cells exhibited innate immunological memory, which could not be reverted by 2-AA re-exposure. Similar responsiveness patterns of ATP and acetyl-CoA as with BMDM cells were observed in murine macrophage RAW 264.7 cells (Figs. S1A and S1B) or human monocyte THP-1 cells (Figs. S1C and S1D) following exposure or re-exposure to 2-AA. These findings further support our observations with BMDM cells and confirm that either cell type can be used to study responses to 2-AA. These results reinforce the notion that 2-AA impacts crucial energy metabolites in macrophages.

**Figure 1.**
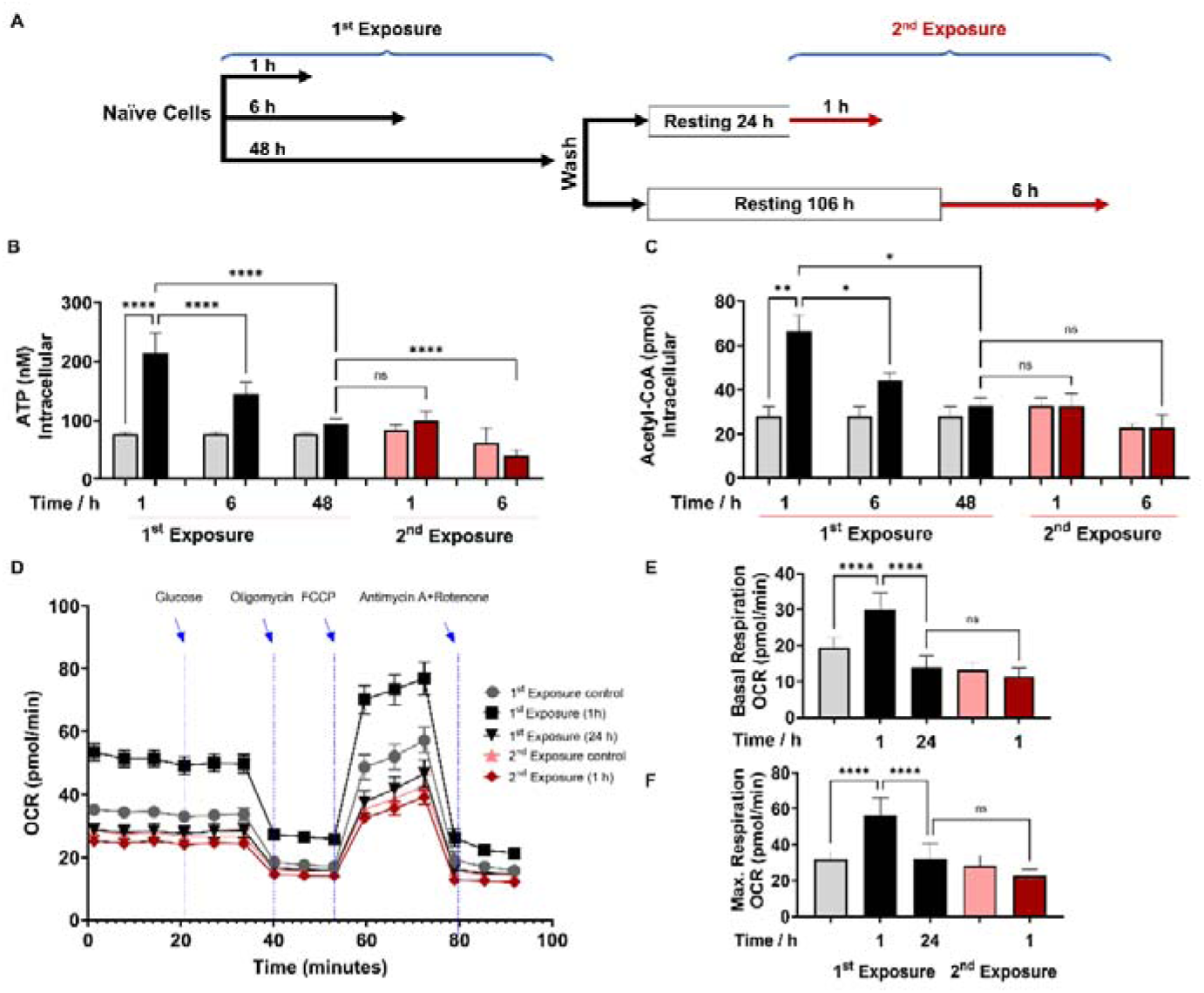
2-AA tolerization decreases crucial metabolites of cellular energy and affects mitochondrial respiration in mouse BMDM. (A) Schematic representation showing experimental design: naïve cells were exposed to 2-AA for 1, 6, or 48 h (black). Cells exposed to 2-AA (400 µM) for 48 h were washed, rested for 24 or 106 h, and re-exposed (200 µM) for 1 or 6 h (red), respectively. The same color code, black cells after 1^st^ exposure and red cells after 2nd exposure, was kept throughout the manuscript with corresponding controls in grey and pink, respectively. The levels of (B) ATP and (C) acetyl-CoA in BMDM cells after 1^st^ and 2^nd^ 2-AA exposure. (D) Real-time oxygen consumption rate (OCR) traces were recorded using a Seahorse XF analyzer and normalized to protein content. Cells were exposed to 2-AA for 1 or 24 h (black), washed and rested for 24 h and re-exposed for 1 h (red). Mitochondrial respiratory parameters, (E) basal respiration, and (F) maximal respiration. Data are presented as mean ± SD, n ≥ 4, \**p* < 0.05, \*\**p* < 0.01, \*\*\**p* < 0.001, and ns indicates no significant difference. One-way ANOVA followed by Tukey’s post hoc test was applied.

Moreover, we sought to determine whether 2-AA-tolerized BMDM cells would respond to heterologous stimulus. To test this, we re-exposed the cells to lipopolysaccharide (LPS), an outer membrane bacterial component that strongly induces macrophages. LPS re-exposure of the 2-AA tolerized BMDM cells did not significantly (*p* = 0.9) increase ATP levels (Fig. S2), indicating that 2-AA cross-tolerized the cells to this heterologous bacterial immunostimulant. On the other hand, as expected, we observed increased ATP levels following LPS stimulation of the non-tolerized BMDM cells (Fig. S2). The broad spectrum of 2-AA cross-tolerization strongly suggests an interplay of epigenetic mechanism rather than ligand-receptor mediated signaling-based mechanism in bringing out the observed 2-AA-mediated tolerance.

### Tolerization by 2-AA leads to a quiescent state by reducing cell bioenergetics and compromising OXPHOS

To further investigate the 2-AA’s impact on the bioenergetics of macrophages, we assessed by Seahorse assay the oxygen consumption rate (OCR) as an index of OXPHOS in 2-AA exposed (for 1 and 24 h) and re-exposed (2^nd^ exposure for 1 h) BMDM cells (Fig. 1D). Values of basal (Fig. 1E) and maximal (Fig. 1F) mitochondrial respiration and spare respiratory capacity (Fig. S3) interpreted from these measurements revealed that 1 h 2-AA exposure of naïve cells significantly increased OCR compared to naïve control cells. At 24 h post-2-AA exposure, cells had a significantly reduced basal OCR level than the control BMDM cells, indicating decreased OXPHOS and an overall quiescent phenotype of tolerized macrophages. Re-exposure of tolerized macrophages with 2-AA also did not augment maximal and basal OCR (Figs. 1D-1F), indicating the unresponsiveness of tolerized BMDM cells. Ultramicroscopic examination of RAW 264.7 cells exposed to 2-AA for 48 h although shows the same number of mitochondria as in control cells, their mitochondrial morphology is altered, appearing smaller and round and having reduced cristae indicating dysfunctional mitochondria (Fig. S4).

These OCR data are consistent with previous direct measurements of ATP (Fig. 1B) and acetyl-CoA (Fig. 1C) and support the notion that 2-AA tolerization induces a quiescent phenotype in these cells characterized by defective OXPHOS. These findings confirm that 2-AA mediates energy homeostatic and metabolic alterations in the immune cells and that 2-AA tolerization arrests these cells in a sustained 2-AA-unresponsive state.

### The pyruvate transport into mitochondria is decreased in tolerized macrophages

Since acetyl-CoA and ATP levels were reduced due to 2-AA tolerization, we investigated whether the mitochondrial dysfunction observed is related to pyruvate metabolism. Pyruvate links glycolysis with mitochondrial production of acetyl-CoA and ATP *via* the tricarboxylic acid (TCA) cycle and OXPHOS [7]. RAW 264.7 cells initially exposed to 2-AA for 1 or 3 h (black bars) exhibited significantly higher concentrations of cytosolic and mitochondrial pyruvate compared to the corresponding naïve control cells (Fig. 2A). Pyruvate significantly decreased over time in cytosolic and mitochondrial fractions as cells were entering into the tolerized state (Fig. 2A). Tolerized cells remain unresponsive to 2-AA re-exposure (red bars), exhibiting no significant change in the concentration of pyruvate in the cytosolic or mitochondrial fraction (Fig. 2A). A similar result was observed in THP-1 cells (Figs. S1E and S1F).

**Figure 2.**
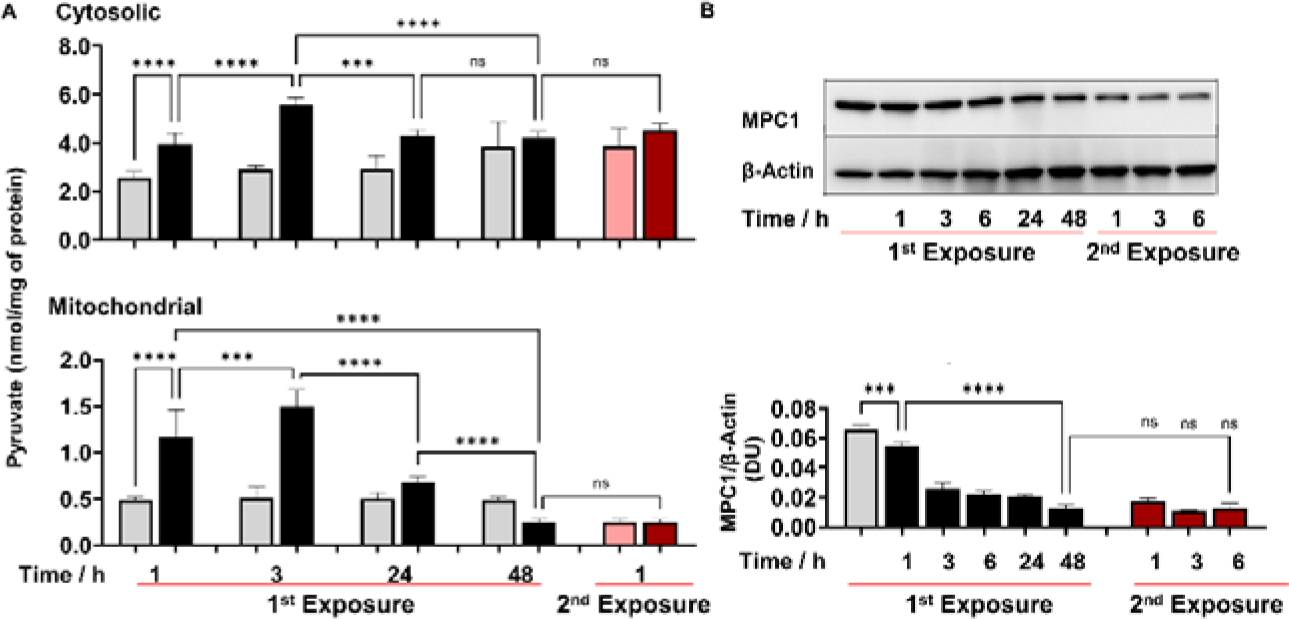
2-AA perturbs the mitochondrial MPC1-mediated import and metabolism of pyruvate. (A) Cytosolic and mitochondrial pyruvate levels following 2-AA exposure (black) or re-exposure (red) and corresponding controls in grey and pink, respectively, for indicated time points. (B) Representative Western blot and results of densitometric analysis of MPC1 protein levels following 2-AA exposure or re-exposure for indicated time points. β-actin was used as a control. Corresponding controls are shown in grey or pink, respectively. Mean ± SD is shown, *n* = 3, \*\*\**p* < 0.001, \*\*\*\**p* < 0.0001, and ns indicates no significant difference. One-way ANOVA followed by Tukey’s post hoc test was applied.

To gain a better understanding of the 2-AA-mediated reductions in mitochondrial pyruvate, ATP production, and acetyl-CoA levels, we sought to determine the involvement of the MPC1 [43] that is crucial in transporting pyruvate from the cytosol into the mitochondria [44]. Western blot studies of whole cell lysates of RAW 246.7 cells exposed to 2-AA showed a significant decrease over time in the abundance of MPC1 protein, and tolerized cells remain unresponsive to 2-AA re-exposure (Fig. 2B). Together, these findings suggest that 2-AA tolerization dysregulates MPC1-mediated transport of the pyruvate into mitochondria.

### 2-AA tolerization suppresses ERRα binding to the *MPC1* promoter and the interaction between PGC-1α and ERRα

Given that the PGC-1α/ERRα axis is the transcriptional regulatory node that activates the expression of MPC1 [33], we determined the protein levels of ERRα. Western blot studies of whole cell lysates showed a significant decrease over time in the abundance of ERRα protein following 1^st^ exposure of RAW 246.7 cells to 2-AA, while tolerized cells remain unresponsive to 2-AA re-exposure (Fig. 3A). By assessing the cytosolic and nuclear abundance of ERRα in tolerized cells we found that both fractions had lower levels of ERRα protein than their corresponding control cells (Fig. 3B).

**Figure 3:**
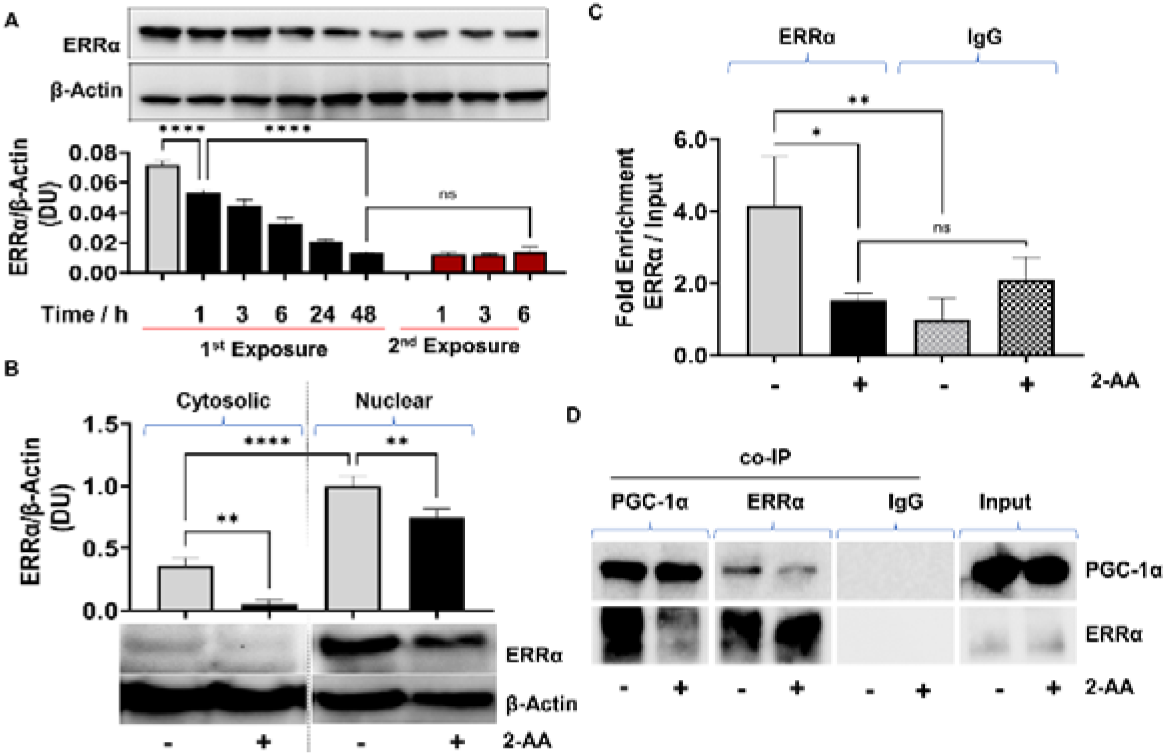
2-AA-mediated macrophage tolerization deranges PGC-1α/ERRα-dependent metabolic programing. (A) Representative Western blot and results of densitometric analysis of ERRα protein levels following 2-AA exposure or re-exposure for indicated time points. β-actin was used as a control. (B) Western blots and its corresponding densitometric analysis of ERRα in cytoplasmic or nuclear lysates of tolerized macrophages exposed to 2-AA for 24 h or not exposed to 2-AA. (C) ChIP-qPCR assay of ERRα binding at the MPC1 promoter in RAW 264.7 tolerized macrophages exposed to 2-AA (200 μM) for 24 h (black) compared to untreated control macrophages (grey). IgG served as a negative control (D) Representative Western blot of co-immunoprecipitation (co-IP) studies of ERRα and PGC-1α in nuclear extracts of 2-AA tolerized (24 h) and control RAW 264.7 cells. Pull down with IgG served as a negative control. 2-AA tolerized macrophages are shown in black, and untreated control macrophages in grey. Mean ± SD is shown, n ≥ 3, \**p* < 0.05, \*\**p* < 0.01, \*\*\*\**p* < 0.0001, and ns indicates no significant difference. One-way ANOVA followed by Tukey’s post hoc test was applied.

Furthermore, because an ERRα binding putative DNA motif (TNAAGGTCA) has been identified upstream of the *MPC1* promoter in humans [43], and we found the same motif to be present 1.5 kB upstream of the transcription start site of mouse *MPC1* promoter, we analyzed if tolerization led to reduced ERRα binding to the *MPC1* promoter motif TNAAGGTCA. Chromatin immunoprecipitation (ChIP) assay followed by qPCR analysis revealed that in tolerized cells, ERRα binds approximately 4-fold less efficiently to the putative binding site on the *MPC1* than in corresponding control cells (Fig. 3C).

Since the transcriptional activity of ERRα is regulated through a protein-protein interaction with the transcriptional coactivator PGC-1α [33], we assessed whether the formation of the PGC-1α/ERRα complex is affected in 2-AA tolerized cells. Using paraformaldehyde fixed nuclear lysates of RAW 264.7 tolerized and naïve cells, we performed a co-immunoprecipitation (co-IP) of PGC-1α and reversed co-IP with ERRα followed by immunoblotting (Fig. 3D). As shown in Figure 3D less ERRα was detected when PGC-1α was pulled down in presence of 2-AA than in naïve control cells. Moreover, reverse co-IP with ERRα protein showed lower levels of PGC-1α in the presence of 2-AA than in naïve control cells (Fig. 3D). Control IgG showed no detectable levels of ERRα following nuclear fraction pull down. These results indicate that in the tolerized macrophages, PGC-1α/ERRα interaction is impaired (Fig. 3D).

Moreover, since PGC-1α enhances the transcriptional activity of ERRα through their interaction, and ERRα transcription is regulated *via* an autoregulatory loop [34], we examined the effect of tolerization on the transcription of *PGC-1*α*, ERR*α, and its target gene *MPC1*. Using RAW 264.7 cells infected with the wild type *PA* strain PA14 or isogenic Δ*mvfR* mutant, which does not produce 2-AA [21] we observed lower RNA transcript levels of *MPC1* (Fig. 4A), *ERR*α (Fig. 4B) and *PGC-1*α (4C) in PA14 compared to Δ*mvfR*. The addition of 2-AA to Δ*mvfR* decreased the levels of these gene transcripts to levels similar to the PA14 infection condition, supporting the role of 2-AA in effect. To test if the observed effects are due to MPC1-dependent reduction of mitochondrial ATP generation, we supplemented the macrophages with ATP. The findings indicate that the addition of ATP in PA14-infected cells elevated the transcript levels of *ERR*α, *MPC1*, and *PGC-1*α, reaching levels similar to those observed in Δ*mvfR* infected macrophages (Fig. 4A, 4B, and 4C). These results indicate that reduced MPC1 function is due to 2-AA tolerization on the transcriptional activation of *MPC1* through the PGC-1α/ERRα axis.

**Figure 4:**
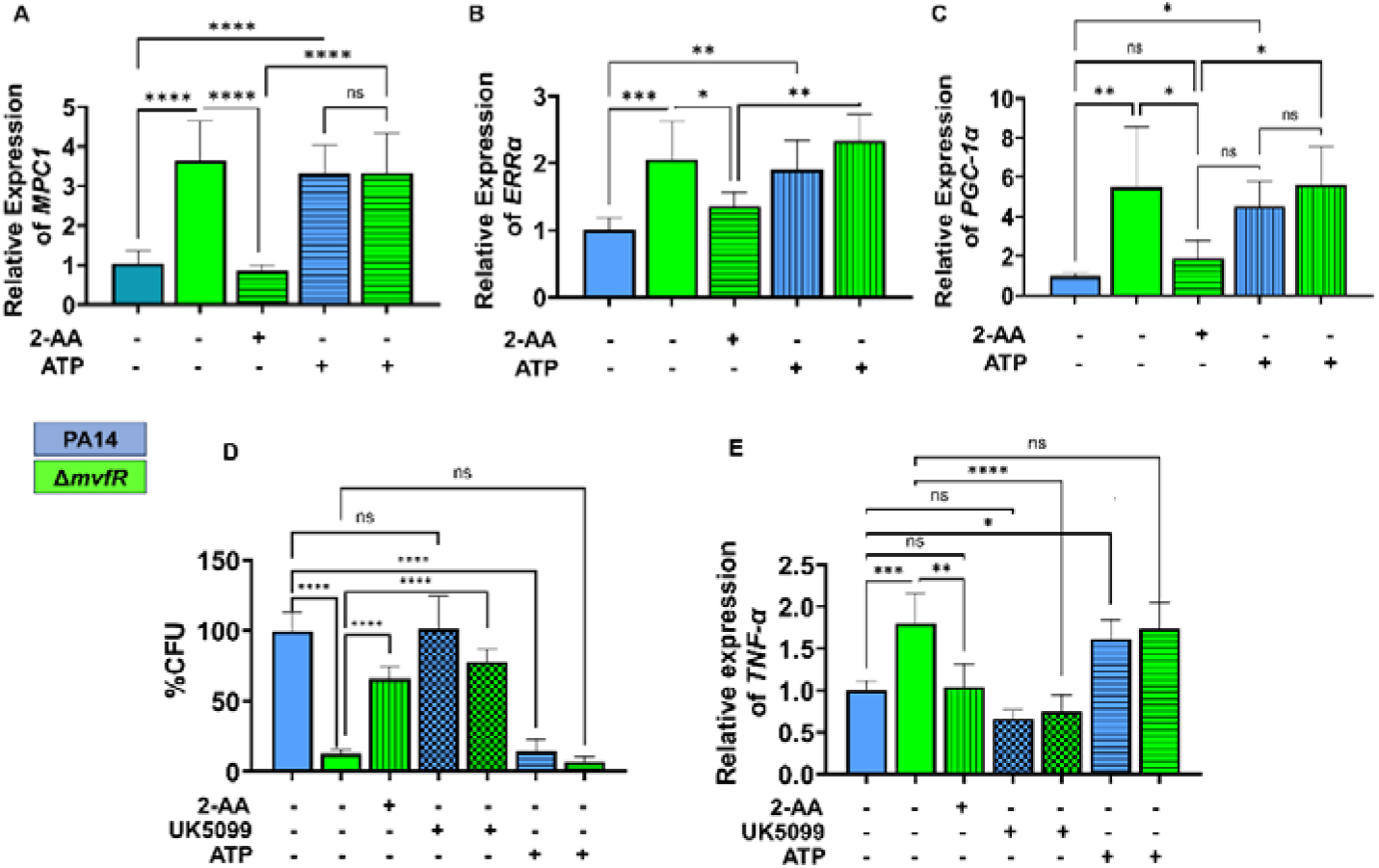
Increased intracellular burden in macrophages is associated with decreased expression of *MPC1, ERR-*α, and *TNF-*α *genes*. Real-time PCR analysis of *MPC1* (A), *ERR-*α (B), and *PGC-1*α (C) expression in RAW 246.7 macrophages infected with PA14 or Δ*mvfR* in the presence or absence of exogenous addition of 2-AA or ATP for 6 h as indicated. Transcript levels were normalized to 18srRNA. PA14-infected cells served as controls. (D) The intracellular burden of PA14 or Δ*mvfR* of infected macrophages in the presence or absence of exogenous addition of 2-AA, UK5099, or ATP. Untreated cells infected with PA14 were set as 100%. (E) Real-time PCR analysis of TNF-α expression in RAW 246.7 macrophages infected with PA14 or Δ*mvfR* in the presence or absence of exogenous addition of 2-AA, ATP, or UK5099. Transcript levels were normalized to 18srRNA. PA14-infected cells served as controls. The compound concentration used for UK5099 was 10 µM and ATP 20 µM. Mean ± SD is shown, *n* = 3, * *p* < 0.05, \*\**p* < 0.01, \*\*\**p* < 0.001, \*\*\*\**p* < 0.0001***, and ns indicates no significant difference. One-way ANOVA followed by Tukey’s post hoc test was applied.

### 2-AA tolerization impairs macrophage-mediated intracellular bacterial clearance through decrease in MPC1-mediated pyruvate import, ATP, and TNF-α levels

2-AA-mediates persistence of *PA in vivo,* dampens proinflammatory responses, and increases the intracellular burden of this pathogen in macrophages *via* epigenetic modifications [17, 18, 26]. Here, we used RAW 264.7 cells infected with the PA14 or the 2-AA deficient Δ*mvfR* mutant to assess the clearance of *PA* by macrophages. Macrophages infected with PA14 showed increased bacterial burden than the cells infected with Δ*mvfR,* and 2-AA addition to Δ*mvfR* led to increased bacterial burden (Fig. 4D). Furthermore, we used UK5099 and ATP to interrogate whether MPC1 mediated mitochondrial pyruvate import and bioenergetics are linked to the clearance of *PA* intracellular burden. The addition of the UK5099 inhibitor strongly enhanced the bacterial intracellular burden in Δ*mvfR* infected macrophages compared to the non-inhibited Δ*mvfR* infected cells, reaching a similar burden to those infected with PA14 (Fig. 4D). Conversely, exogenously added ATP to macrophages infected with PA14 strongly reduced the *PA* intracellular burden (Fig. 4D).

Given that 2-AA tolerization decreases the expression of pro-inflammatory cytokine TNF-α by hypo-acetylating the core histone 3 lysine 18 acetylation (H3K18ac) mark at TNF-α promoter [18], we investigated the link between bioenergetics and TNF-α expression in infected macrophages. UK5099, or ATP was added exogenously to infected macrophages with PA14 or Δ*mvfR* (Fig. 4E). As shown in Figure 4E, PA14 infected cells showed lower TNF-α transcript levels compared to the Δ*mvfR* infected cells. Supplementation of 2-AA to Δ*mvfR* infected cells led to a decrease in the TNF-α transcript levels (Fig. 4E). The addition of the MPC1 inhibitor, UK5099 in Δ*mvfR* infected cells counteracted the increase in TNF-α transcript levels observed in Δ*mvfR* infected cells in the absence of UK5099 (Fig. 4E). Conversely, exogeneous addition of ATP to PA14 infected cells enhanced the transcription of TNF-α compared to the untreated PA14 infected cells, while no difference in TNF-α expression levels were observed in Δ*mvfR* infected macrophages in presence or absence of ATP (Fig. 4E).These findings strongly suggest that 2-AA tolerization severely alters macrophages’ ability to facilitate the clearance of *PA* intracellular burden, *via* the reduction in MPC1-mediated pyruvate import into mitochondria, ATP levels and TNF-α transcription.

### *In vivo* studies corroborate the 2-AA-mediated decrease *in vitro* of the central metabolic fuel acetyl-CoA and the energy-carrying molecule ATP and their association with *PA* persistence

To determine whether the decrease in the key metabolites mediated by 2-AA is also observed *in vivo* during infection, we quantified the levels of ATP and acetyl-CoA in murine spleen tissues at 1, 5, and 10 days (Fig. 5). This organ was selected for our immunometabolic studies due to its role in regulating not only local but also systemic (whole-body) immune responses, facilitated by various immune cells including macrophages [45]. Mice were infected with *PA* strain PA14, or isogenic mutant Δ*mvfR,* which does not produce 2-AA [21]. It is important to note that it is not possible to generate or utilize a bacterial mutant that is defective in 2-AA only because 2-AA is formed by spontaneous decarboxylation rather than by an enzyme-catalyzed reaction [46–48]. Therefore, animals that were either infected with Δ*mvfR* and received 2-AA (Δ*mvfR+*2-AA) at the time of infection or uninfected mice injected with a single dose of 2-AA served as direct controls (Fig. 5). Naïve and sham mice groups served as additional basal controls (Fig. 5).

**Figure 5.**
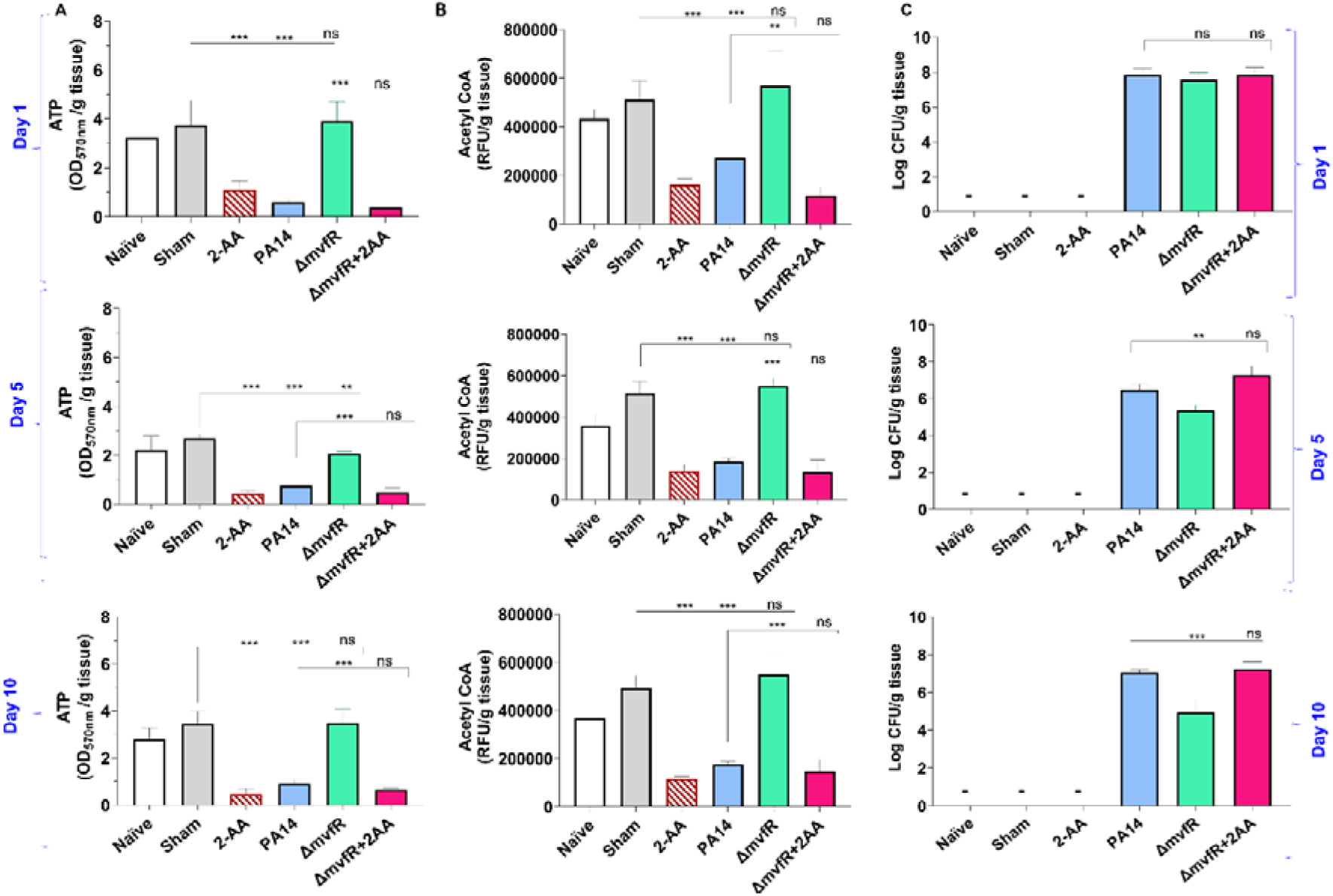
2-AA promotes a long-lasting decrease in ATP, acetyl-CoA levels, and bacterial persistence in *PA*-infected mice. (A) ATP and (B) acetyl-CoA concentrations in the spleens of mice infected with *PA* wild type (PA14), the isogenic mutant Δ*mvfR*, Δ*mvfR* injected with 2-AA at the time of infection (Δ*mvfR* + 2-AA), or non-infected but injected with 2-AA (6.75 mg/kg), (C) Bacterial burden in muscles expressed as CFU count was analyzed using the Kruskal–Wallis non-parametric test with Dunn’s post-test; \*\*\**p* < 0.001, and ns indicates no significant difference. Control mice groups: naïve were not given 2-AA; mice receiving 2-AA were given a single intraperitoneal injection of 2-AA; sham represents a burn/PBS group since the burn and infection model was used. Results of three independent replicates with 4 mice per group for 1, 5, and 10 d are shown. Means ± SD are shown, \**p* < 0.05, \*\**p* < 0.01, \*\*\**p* < 0.001, and ns indicate no significant difference. One-way ANOVA followed by Tukey’s post hoc test was applied.

Infection with PA14 or uninfected mice injected with 2-AA led to a significant decrease in both ATP (Fig. 5A) and acetyl-CoA (Fig. 5B) concentrations in spleen tissues compared to Δ*mvfR* infected mice that sustained higher ATP and acetyl-CoA levels similar to naïve and sham control groups across the time points tested. However, in mice infected with Δ*mvfR* and 2-AA injected (Δ*mvfR*+2-AA) at the time of infection, ATP and acetyl-CoA concentrations in the spleen decreased to levels comparable to those of PA14-infected animals, strongly indicating the biological function of 2-AA in decreasing these key metabolites. These *in vivo* findings further support the adverse action of 2-AA on host energy homeostasis and metabolism observed in *in vitro* studies.

The metabolic alterations observed are associated with the host tolerance to *PA* persistence (Fig. 5C). Using *Pseudomonas* isolation agar plates, we evaluated the bacterial load at the infection site over the course of 10 days, obtaining samples at 1, 5, and 10 days. At one day post-infection, mice infected with PA14, Δ*mvfR,* or Δ*mvfR*+2-AA exhibited no difference in bacterial burden (Fig. 5C), verifying the ability of all strains to establish infection. At 5- and 10 days post-infection, mice infected with PA14 or Δ*mvfR*+2-AA sustained the bacterial burden at a significantly higher bacterial burden over time compared to those infected with Δ*mvfR* (Fig. 5C).

To strengthen the relevance of our *in vivo* data, we performed additional *in vivo* experiments. In this set of *in vivo* studies, mice received the first exposure to 2-AA by injecting 2-AA only and the 2^nd^ exposure through infection with PA14 or Δ*mvfR* four days post-2-AA injection. As shown in the supplementary Figure S5, the levels of ATP and acetyl CoA in the spleen of infected animals and the enumeration of the bacterial counts were similar between PA14 or Δ*mvfR* receiving the 1^st^ 2-AA exposure and agree with the “one-shot infection” findings presented in Figure 5 with the PA14 or Δ*mvfR* +2-AA infected mice or those receiving 2-AA only. These results are consistent with our previous findings, showing that 2-AA impedes the clearance of PA14 [17, 18] and provide compelling evidence that the metabolic alterations identified may favor *PA* persistence in infected tissues.

## Discussion

The *PA* signaling molecule 2-AA that is abundantly produced and secreted in human tissues [21, 22] is the first QS molecule that epigenetically reprograms immune functions, promotes immune tolerance, and sustains *PA’s* presence in host tissues [17, 18, 22, 26]. The present study provides insights into the mechanistic actions of 2-AA on cellular metabolism that contribute to host immune tolerance to *PA* persistence. We uncover that this signaling molecule causes distinct metabolic alterations in macrophages’ mitochondrial respiration and energy production promoted via the PGC-1α/ERRα axis and MPC1-dependent OXPHOS that links the TCA cycle to the production of ATP and energy homeostasis (Fig. 6). Our results show that 2-AA tolerization decreases ATP the crucial energy metabolite and the histone acetylation metabolite acetyl-CoA and *in vivo*, implicating the importance of the key energy-producing mitochondrial process in immune tolerance. Although macrophages respond to the first exposure to 2-AA by enhancing ATP and acetyl-CoA production, long-term exposure to 2-AA leads to a tolerized state characterized by sustained reduced ATP and acetyl-CoA concentrations, a quiescent bioenergetic state with reduced mitochondrial pyruvate uptake and unresponsiveness to a 2^nd^ exposure.

**Figure 6:**
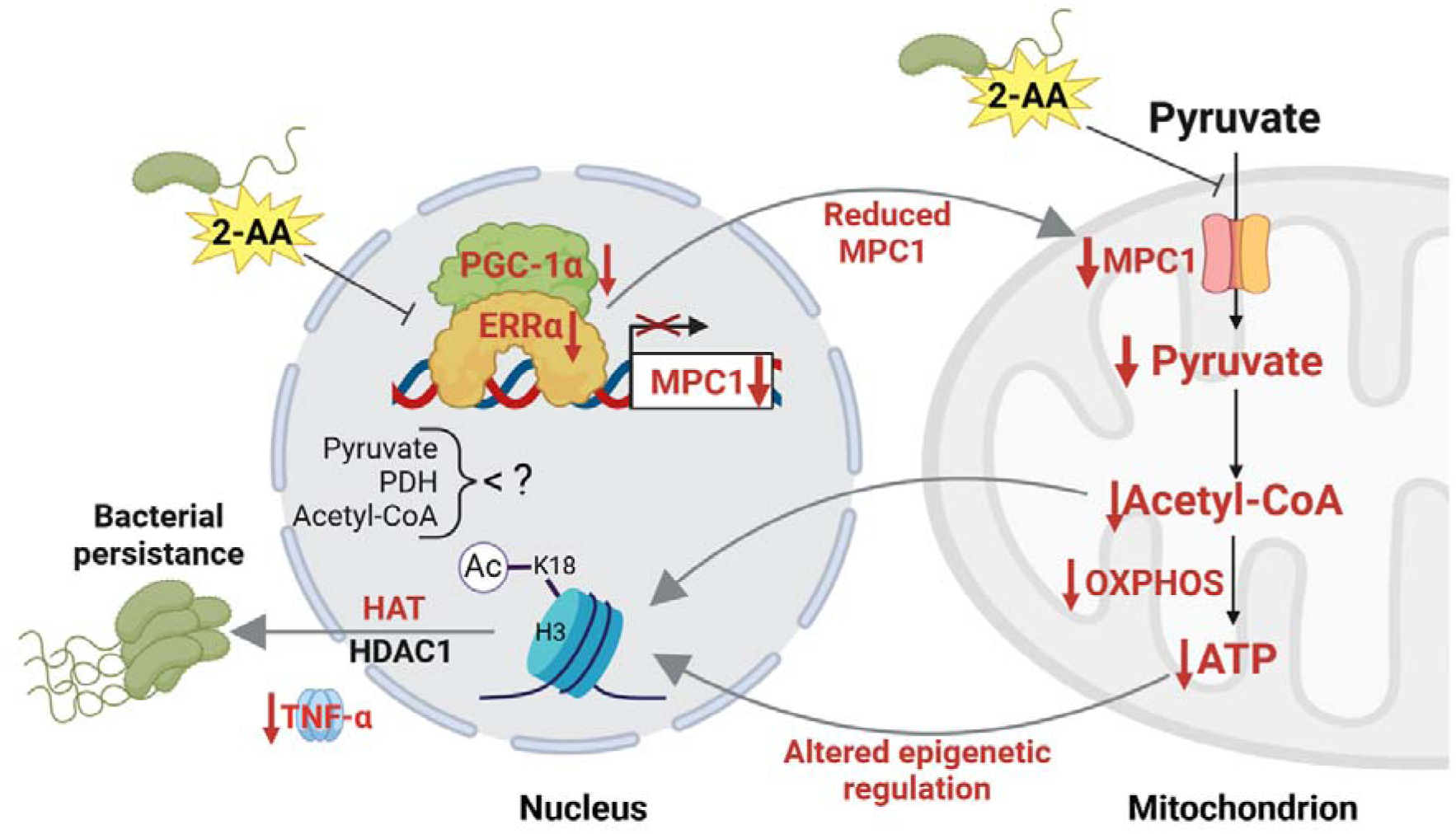
Proposed mechanism by which 2-AA impairs bioenergetics through the inhibition of MPC1-mediated pyruvate transport into mitochondria and its impact on the PGC-1α/ERRα axis. 2-AA tolerized macrophages exhibit diminished pyruvate levels in mitochondria due to the decreased expression of *MPC1*, a consequence of the 2-AA impact on the interaction of ERRα with the transcriptional coactivator PGC-1α for the effective transcription of ERRα since ERRα controls its own transcription and that of MPC1. In the presence of 2-AA, the weakened interaction between ERRα and PGC-1α results in reduced expression of MPC1 and ERRα. The reduction in mitochondrial pyruvate levels leads to decreased acetyl-CoA and ATP levels, which modulate HDAC1- and HAT-catalyzed remodeling of H3K18 acetylation. The diminished levels of this epigenetic mark have previously been associated with an increased intracellular presence of bacteria in macrophages, as demonstrated by our group [18]. Pathways, proteins, and metabolites that are negatively affected are indicated in red, while positively affected are denoted in black. The figure was created using BioRender.com.

Our results reveal that the decrease of ATP and acetyl-CoA in tolerized macrophages results from the 2-AA-mediated perturbation of the physical interaction between ERRα and PGC-1α. The PGC-1α/ERRα axis has not previously been implicated in tolerization but has been extensively studied in cancer, underscoring the novelty of our findings. Studies in cancer cells have shown that the physical interaction of human ERRα with PGC-1α strongly enhances the binding of ERRα to DNA to induce the activation of the targeted genes [33], including *MPC1,* responsible for the import of cytosolic pyruvate into mitochondria and PDH, that catalyzes the conversion of pyruvate to acetyl-CoA in mitochondria. Inhibition of ERRα has also been shown in cancer cell metabolism studies to interfere with pyruvate entry in mitochondria by inhibiting the expression of MPC1 [49], and inhibition of ERRα led to decreased expression of PGC-1α-controlled expression of mitochondrial genes in mice brains [45]. Indeed, we find that the transcription of *ERR*α*, PGC-1*α, and *MPC1* genes and pyruvate uptake into mitochondria is reduced in tolerized macrophages. Given that pyruvate is a primary carbon source in the TCA cycle that fuels OXPHOS and the production of ATP into mitochondria, it explains the reduced ATP and acetyl-CoA levels observed in tolerized macrophages. Future studies will focus on unveiling the pathways by which macrophages respond to the bioenergetic changes identified and decipher their effect on the pyruvate cycle in host tolerance to infection [50–52].

We show that the unprecedented action of a QS bacterial small signaling molecule on the interaction between ERRα and PGC-1α in the nucleus is hampered. Although it remains to be elucidated whether 2-AA interferes directly or indirectly with the PGC-1α/ERRα axis, our data bring up the possibility that the reduced production of acetyl-CoA may be responsible since cholesterol is a known coactivator/ligand of ERRα [53], is generated from acetyl-CoA [53]. This possibility may also explain the reduced expression of *ERR*α and lower abundance since cholesterol is also required for its activation and autoregulation. However, it is also possible that 2-AA may antagonize the interaction between ERRα and PGC-1α by binding to ERRα. Small molecules as antagonists of ERRα have been reported previously [54]. The weaker binding of ERRα to the *MPC1* promoter site, and consequently reduced MPC1 expression, explains the reduced pyruvate presence in mitochondria and that of ATP and mitochondrial quiescence. These findings open avenues for exploring novel therapeutic strategies through the overexpression of ERRα to counteract the tolerization induced by 2-AA and enhance clearance of *PA* infection. Interestingly, ERRα overexpression on breast cancer metastases promotes an efficient antitumor immune response selectively in the bone [55].

The observed cellular metabolic perturbances also align with our previous studies in skeletal muscle, which pointed to mitochondrial dysfunction triggered by 2-AA [24, 25]. Interestingly, the *PA* QS LasR-regulated homoserine lactone molecule, 3-oxo-C12-HSL, was reported to attenuate the expression of PGC-1α and decrease the mitochondrial respiratory capacity in lung epithelial cells [27]. Moreover, another *PA* QS molecule, PQS, also regulated by LasR, was recently shown to induce organelle stress, including mitochondria, by disrupting the mitochondrial membrane potential in human macrophages [56]. As opposed to 2-AA, which dampens the pro-inflammatory response, PQS increases proinflammatory cytokines [56], consistent with the fact that PQS is an acute infection type molecule unrelated to chronic/persistent infections.

Previously, we reported that 2-AA tolerization induces histone deacetylation via HDAC1, reducing H3K18ac at the TNF-α promoter [18]. The findings with acetyl-CoA reduction, the primary substrate of histone acetylation, and the TNF-α transcription using UK5099 and ATP in 2-AA treated macrophages are in support of the bioenergetics disturbances observed in macrophages and their link to epigenetic modifications we have shown to be promoted by 2-AA [18]. Macrophages exposed to 2-AA in the presence of exogenous ATP showed improved intracellular bacterial clearance and enhanced TNF-α levels, supporting the epigenetic interconnection of macrophages and cellular ATP responsiveness levels against infection. The exogenous addition of UK5099 that reverted the efficiency of the *mvfR* infected macrophages to clear the *PA* intracellular burden and the counteraction to the 2-AA effect by ATP addition that increases the clearance of *PA* intracellular burden support the importance of this energy-carrying molecule in *PA* persistence.

The results with the MPC1 inhibitor, UK5099, suggest that the availability of pyruvate could underlie the mechanism of 2-AA-regulated HDAC/HAT-dependent control of transcription of pro-inflammatory mediators and bacterial clearance [57]. These results align with our previous findings, showing that establishing bacterial intracellular burden in macrophages is HDAC1 dependent [57]. It remains to be elucidated if HDAC/HAT-mediated histone modification directly regulates the expression of OXPHOS genes in 2-AA tolerized macrophages, as it was shown that histone deacetylation downregulates OXPHOS in persistent *M. tuberculosis* infection [58, 59]. Given the complexity of 2-AA-mediated long-term effects, future omics studies combined with immune profiling may aid in deciphering the possible network of co-factors across different subpopulations of immune cells and the immunometabolic reprogramming related to 2-AA.

We have shown that although 2-AA tolerization leads to a remarkable increase in the survival rate of infected mice, it permits an HDAC1-dependent sustained presence of *PA* in mice tissues and intracellularly in macrophages [17, 18, 22, 26]. Here, we use 2-AA-producing and isogenic non-producing *PA* strains to confirm the 2-AA-mediated decrease in the central metabolite acetyl-CoA and the energy-carrying molecule ATP during infection. These murine infection studies provide strong evidence of the 2-AA biological relevance in reducing these metabolites *in vivo*. Taking together our previous and current *in vivo* findings provide compelling evidence that the metabolic alterations identified favor *PA* persistence in infected tissues.

That 2-AA permits *PA* to persist in infected tissues despite rescuing the survival of infected mice underscores its difference from that of LPS tolerization, which results in bacterial clearance [23] *via* the mechanism that relies on a set of different HDAC enzymes. LPS tolerization predominantly involves changes in H3K27 acetylation [60], while 2-AA tolerization involves H3K18 modifications [22]. The 2-AA-mediated effects reported here are also distinct from the epigenetic-metabolic reprogramming mediated by LPS [61], which promotes endotoxin tolerance in immune cells by upregulating glycolysis and suppressing OXPHOS [62]. Although 2-AA and LPS implicate different components and lead to different outcomes, both involve epigenetic mechanisms and immune memory.

Metabolic reprogramming upon infection may be pathogen-specific, with each pathogen impacting specific metabolic pathways that better fit its respective metabolic needs [7]. This was shown for infections of various tissues with *Chlamydia pneumonia*, *Legionella pneumophila*, *Mycobacterium tuberculosis*, and *Salmonella* [63–66] and, by our group, for *PA* infections [17, 18, 24]. It would be interesting, however, to test whether 2-AA affects the killing efficacy of macrophages against other pathogens, as emerging evidence shows that synergistic or antagonistic interactions between clinically relevant microorganisms and host have important implications for polymicrobial infectious diseases [67].

While in this study, we focused on the role of ERRα mainly in pyruvate metabolism; future studies are needed to reveal other ERRα/PGC-1α axis-dependent metabolic and immune pathways are modulated by 2-AA. In this view, the cholesterol synthesis pathway that relies on acetyl-CoA precursor, as cholesterol is a known coactivator/ligand of ERRα [53] and fatty acid catabolism, generating acetyl-CoA, and PGC-1α regulates fatty acid oxidation [68].

This study unveils the unprecedented actions of a QS bacterial molecule in orchestrating cellular metabolic reprogramming in addition to the epigenetic reprogramming reported previously and has also been shown to promote a long-lasting presence of *PA* in the host [17, 18, 26]. That 2-AA cross-tolerized macrophages to LPS corroborates our previous findings on the implication of epigenetic mechanism [18] rather than ligand-receptor mediated signaling-based mechanism and raises the possibility that this QS molecule may confer non-specific cross-protection. This is an aspect we plan to investigate in the future.

The reported immunometabolic reprogramming contributes to a better understanding of the molecular and cellular mechanisms, biomarkers, and functional significance that may be involved in immune tolerance to persistent infection, providing for designing and developing innovative therapeutics and interventions. These approaches can focus on promoting host resilience against bacterial burden and safeguarding patients from recalcitrant persistent infections.

## Materials & Methods

### Ethics statement

The animal protocol was approved by the Institutional Animal-Care and Use Committee (IACUC) of Massachusetts General Hospital (protocol no. 2006N000093). No randomization or exclusion of data points was applied.

### Conditions used for the exposures of the cells to 2-AA

Mouse BMDM cells were isolated from the femur of CD1 6 weeks old male and female mice and used to estimate the levels of various metabolites (Fig. 2A) following exposure and re-exposure to 2-AA (Fig. 1A). The 1^st^ exposure of naïve BMDMs (10^6^/mL in 6-well plates) was achieved with 400 µM 2-AA for 1, 6, or 48 h. Cells exposed for 48 h were used for the 2^nd^ 2-AA exposure. These cells were washed to remove residual 2-AA, left in the medium in the absence of 2-AA for either 24 or 106 h, and re-exposed to 2-AA (200 µM) for 1 or 6 h, respectively. For RAW 264.7 (10^6^/mL in 6-well plates) and THP-1 cells (10^6^/mL in 6-well plates), the 2-AA concentration and conditions used were the same as those used for BMDM cells. The hours of the 1^st^ round and 2^nd^ round of exposure to 2-AA are indicated in Figure S1.

For the LPS stimulation studies, cells were stimulated with 400 µM 2-AA or 100 ng/mL LPS for 6 h and 800 µM of 2-AA for 48 h. Cells receiving 1^st^ exposure for 48 h were washed and re-exposed to 400 µM of 2-AA or 100 ng/mL LPS for 6 h.

### ATP and acetyl-CoA quantifications

The levels of ATP acetyl-CoA, and were assessed in BMDM and RAW 264.7 macrophage cells following exposure to 2-AA at the times indicated and by utilizing the ATP Assay kit (cholorimetric/fluorometric) (#ab83355, Abcam) and the PicoProbe AcCoA assay kit (ab87546, Abcam), respectively, according to the manufacturer’s instruction. Quantifications of ATP and acetyl-CoA were performed in triplicates.

#### ATP

For ATP determination, macrophages were lysed and subsequently centrifuged at 14,000 rpm for 10 min at 4 °C. The supernatants were transferred to a fresh Eppendorf tube. Standards and cell supernatants of 50 µL were added to a 96-well plate suitable for fluorescent analysis (black sides, clear bottom). A reaction mixture containing an ATP converter, probe, buffer, and developer mix was then added to all wells (50 µL) and incubated away from light at room temperature for 30 min. ATP quantification was conducted fluorometrically at 535/587 nm using a plate reader and the following settings: λex□=□535□nm; λem□=□587□nm. Fluorescence was measured using a microplate reader (Tecan Group Ltd, Männedorf, Switzerland)

#### Acetyl-CoA

Acetyl-CoA content was assessed by first deproteinizing total cell fractions of macrophages using the perchloric acid and then centrifuging at 14,000 rpm for 10 min at 4 °C. 50 µL of cells’ supernatant sample CoASH were quenched to correct the background generated by free CoASH. Following the homogenization procedure described above, the samples were diluted with the reaction mix, and fluorescence was quantified using a plate reader and the settings as above with ATP.

### Seahorse Assays

Seahorse analysis was performed according to previously published protocols [69]. Briefly, freshly prepared BMDM cells were reseeded in complete RPMI-1640 cells using Seahorse plates (Agilent cat. no. 103729100) at a density of 5 × 10^4^ cells per well. Cells were exposed to 400 µM 2-AA for 1 h or 24 hours (1^st^ exposure). Cells receiving 1^st^ exposure for 24 h were washed and re-exposed to 200 µM of 2-AA for 1 h. Naïve cells were used as control. Prior to initiating Seahorse measurements, cells were washed, and Seahorse XF DMEM supplemented with 2 mM glutamine (Gibco cat. No. 25030-081) was added to each well. Cells were allowed to stabilize in a 37°C incubator without CO_2_ for 1 h. The Seahorse cartridge was hydrated and calibrated as per the manufacturer’s instructions. The Mitochondrial stress test kit (Agilent cat. no. 10395-100) was used (oligomycin at 1 μM, FCCP (1.5 µM), rotenone (0.5 μM), and antimycin A at (0.5 μM)) with slight modifications according to the published protocol [69] that included injection of 25 mM glucose, and sodium pyruvate (1 μM). All samples N= 4 were run in a Seahorse XFe96 Analyzer, and data were analyzed using Wave and plotted using GraphPad Prism software.

### Isolation of cytosolic and mitochondrial fraction and pyruvate quantification

Pyruvate is produced in the cytosol and is transported into the mitochondria. RAW 264.7 cells were plated at 1 ×□10^6^ to incubate overnight at 37°C in a CO_2_ incubator. Cells were exposed to 2-AA for 1, 3, 24, or 48 h or re-exposed for 1 h as described above. Mitochondria and cytosolic fractions of each group were isolated utilizing the Mitochondria Isolation Kit for cultured cells (#ab110170, Abcam) following the manufacturer’s protocol. Briefly, cells were collected with a cell lifter and pelleted by centrifugation at 1,000 g, frozen, and then thawed to weaken the cell membranes. The cells were resuspended in Reagent A and transferred into a pre-cooled Dounce Homogenizer. The homogenates were centrifuged at 1,000 *g* for 10 min at 4°C and saved as supernatant #1 for the cytosolic fractions. The pellet was resuspended in Reagent B, followed by repeat rupturing and centrifugation. The pellet was collected and resuspended in 500 μL of Reagent C supplemented with Protease Inhibitor cocktails (P8340, Sigma Aldrich). Following separation, a Bradford assay was conducted to determine the protein concentration in each fraction. Pyruvate levels were determined in cellular fractions and mitochondrial fractions by using the Pyruvate Assay Kit (#ab65342, Abcam) according to the manufacturer’s instructions. Briefly, after deproteinization using perchloric acid, the samples were neutralized in ice-cold 2 M KOH. 10 µL samples were incubated with reaction mix and kept on the plate at RT for 30 min in the dark. The absorbance was measured in a microplate reader (Tecan Group Ltd, Männedorf, Switzerland) at 570 nm, and the results were shown in three independent experiments.

### Bacterial strains and growth conditions

The *PA* strain known as Rif^R^ human clinical isolate UCBPP-PA14 (also known as PA14) was used [70]. The bacteria were grown at 37 °C in lysogeny broth (LB) under shaking and aeration or on LB agar plates containing appropriate antibiotics. PA14 and isogenic mutant Δ*mvfR* [19] cultures were grown in LB from a single colony to an optical density of 600 nm (OD_600_) of 1.5, diluted 1:50,000,000 in fresh LB media, and grown overnight to an OD_600_ of 3.0.

### Gentamicin protection assay

RAW 264.7 macrophage cells were plated on 6 well-cell culture-treated plates overnight in pyruvate-free DMEM. After 3 h of incubation with 10 µM UK5099, 20 µM ATP or 400 µM 2-AA, cells were infected with PA14 and isogenic mutant Δ*mvfR* at 5 MOI for 30 min at 37 °C in 5% CO_2_. Unbound bacteria were removed by washing once with cold DMEM medium. Afterward, the cells were incubated with 100 µg/ml of gentamicin for 30 min to eliminate residual extracellular bacteria. The cells were washed with DMEM, transferred to free medium without gentamicin and kept for 3 h at 37°C in 5% CO_2_. The infected cells were scraped after 3 h, centrifuged at 500 × g, and lysed in distilled water. The lysed cells were immediately diluted in PBS and plated on LB agar plates to assess bacterial presence. Bacterial colony-forming units (CFUs) were counted after incubating the plates overnight at 37°C.Untreated cells infected with PA14 were set as 100 % and the reduction of the bacterial load was expressed as %CFU.

### Pharmacological inhibitors and ATP supplementations

For the OXPHOS inhibition assay, RAW 264.7 macrophage cells were treated with UK5099 (10 µM, Sigma-Aldrich, dissolved in DMSO 3 h prior to 1^st^ or 2^nd^ rounds of 2-AA exposure. RAW 264.7 macrophages supplemented with ATP (20 µM, Sigma-Aldrich, dissolved in PBS) received ATP at the time of 1^st^ and 2^nd^ 2-AA exposure.

### Co-Immunoprecipitation assay

For all immunoprecipitation assays, protein A/G agarose Magnetic beads (Pierce) were used after washing in 1X IP buffer. Nuclear lysates from RAW 264.7 tolerized cells exposed to 2-AA for 24 h or non-exposed cells were diluted in 1X IP buffer, and 500 µg of protein was taken for each experiment. The lysates were precleared by using unbound 50 µl protein A/G magnetic beads for 2 h at room temperature on a rotator. Precleared lysates were either incubated with ERRα, PGC-1α or rabbit IgG antibody overnight at 4 °C. 100 µl protein A/G magnetic beads were used to pull down the antibody-protein complex and washed twice with IP buffer to remove unbound proteins. Magnetic beads were then eluted in 2X Laemmli SDS-PAGE loading buffer. Immunoprecipitated ERRα and PGC-1α was detected by Western blot analyses, using conformational specific Anti Rabbit antibody (TruBlot).

### Immunoblotting analysis

Cells were plated at 6×10^5^ cells per well in six-well plates. Cells were washed with PBS and subsequently lysed using RIPA lysis buffer containing 1 mM phenylmethylsulfonyl fluoride (PMSF). 20μg of proteins were separated by electrophoresis on any KD (Kilo Dalton) (Bio-Rad, Cat no. 4569033) SDS-polyacrylamide gel. Proteins were transferred to a 0.2 μm polyvinylidene fluoride membrane (PVDF, Millipore, Billerica, USA) using a Bio-Rad semi-dry instrument. After blocking with 5% BSA in TBS containing 0.1% Tween-20 for 1 h at room temperature, the membranes were incubated with a primary antibody ERR-a (Abcam, #ab76228), MPC1 (D2L91, Cell signaling), and anti-β-actin (cat no: sc-47778) (Santa Cruz Biotechnology) overnight at 4°C. Following washing, the membranes were incubated with an anti-rabbit secondary antibody, and the bands were detected by SuperSignal West Pico Chemiluminescent Substrate (Thermo Scientific) reaction, according to the manufacturer’s instructions. The blots were visualized in the ChemiDOC Imaging system (Bio-Rad Laboratories, Inc., Hercules, CA, USA). The bands were analyzed densitometrically using QuantityOne software (Bio-Rad).

### Chromatin immunoprecipitation (ChIP) and ChIP-qPCR

For ChIP studies, RAW 264.7 macrophage cells exposed to 2-AA for 24 h were cross-linked in 1% (v/v) methanol-free formaldehyde for 10 min and then placed in 0.125 M glycine for 5 min at RT. Using the truChIP High Cell Chromatin Shearing kit (Covaris, United States), cells were prepared for sonication according to the Covaris protocol. Approximately 1□×□10^7^ cells were plated in a 12□mm□×□12□mm tube and subjected to shearing with the Covaris S220 sonicator for 8□min (140 peak power, 5 duty factor, 200 cycles/burst). The Magna ChIP A/G kit (Millipore, United States) was used for the subsequent immunoprecipitations according to the manufacturer’s protocol. Briefly, chromatin from approximately 10^6^ cells was incubated overnight at 4°C with 2□μg of anti-ERRα or anti-PGC-1α (Abcam, USA) ChIP-grade antibody and 20□μL of A/G magnetic beads. The beads were washed serially (5□min each) with low-salt wash buffer, high-salt wash buffer, LiCl wash buffer, and TE buffer from the kit at 4°C. Chromatin was eluted with elution buffer containing Proteinase K at 62°C for 4 h, then incubated at 95°C for 10□min. DNA was isolated by column purification (QIAquick PCR purification kit).

Real-time ChIP-qPCR was performed with the Brilliant II SYBR green super mix (Agilent, USA). Forward (AGTGGTGACCTTGAACTTCCC) and reverse (CTGAAGACGACCTTCCCCTT) primers were chosen to amplify a genomic locus of *MPC1* promoter, which had a putative ERR binding site (TNAAGGTCA) at 1582 bp upstream of the start site. Normalized values were calculated using the percent-input method relative to the IgG. The assay was performed three times.

### RNA extraction and RT-qPCR

Total RNA from all the groups mentioned above was isolated from approximately 2□×□10^6^ cells with the RNeasy minikit (Qiagen, USA), and cDNA was prepared with the iScript Reverse transcription kit (Bio-Rad, USA), as per the manufacturer’s instruction. Real-time PCR was conducted using the PowerUP SYBR Green Master mix (Applied Biosystem, USA) and primer sets for mouse ERR-α (forward: ACTACGGTGTGGCATCCTGTGA; reverse: GGTGATCTCACACTCATTGGAGG), *PGC-1*α (forward: GAATCAAGCCACTACAGACACCG; reverse: CATCCCTCTTGAGCCTTTCGTG), MPC-1 (forward: CTCCAGAGATTATCAGTGGGCG; reverse: GAGCTACTTCGTTTGTTACATGGC), TNF-α (forward: GGTGCCTATGTCTCAGCCTCTT; reverse: GCCATAGAACTGATGAGAGGGAG) and mouse 18S rRNA (forward: GTTCCGACCATAAACGATGCC; reverse: TGGTGGTGCCCTTCCGTCAAT). The transcript levels of all the genes were normalized to 18S rRNA with the ΔΔCT method. The relative expression was calculated by normalizing transcript levels to those of PA14-infected cells. The assay was conducted in triplicate; means and standard deviations were calculated for each group.

### Animal infection and metabolites assessment experiments

The full-thickness thermal burn injury and infection (BI) model [17, 18, 71] was used to assess the effect of 2-AA on metabolic alterations in the spleens of six-week-old CD1 male mice and bacterial burden (Charles River Labs, USA). A full-thickness thermal burn injury involving 5-8% of the total body surface area was produced on the shaved mouse abdomen dermis, and an inoculum of ∼1 × 10^4^ PA14 and Δ*mvfR* [19] cells in 100 μL of MgSO_4_ (10 mM) was injected intradermally into the burn eschar. For the groups that received 2-AA, mice were injected intraperitoneally (IP) with 100 μL of 2-AA (6.75 mg/kg). The entire procedure was done under the influence of anesthesia. One of the groups of mice infected with the PA14 isogenic mutant Δ*mvfR* [19] also received 100 μL of 2-AA in PBS (6.75 mg/kg) at the time of infection (Δ*mvfR* +2-AA) and served as an additional control. CFU counts from rectus abdominus muscle (underlying the burn-infected tissue) were assessed in groups of 4 mice each at 1, 5, and 10 d post-BI by plating diluted muscle homogenate on Pseudomonas isolation agar (Sigma-Aldrich) plates containing rifampicin (50 mg/L). Spleen samples from mice were collected for metabolite analyses from all mice groups at 1, 5, and 10 d post-BI to assess ATP (#ab83355 Abcam) and acetyl-CoA (#ab87546, Abcam) as described above. Tissues were homogenized, 100 mg of homogenate was centrifuged at 4°C for 15 min at 10,000 g, and supernatants were collected. The supernatant was mixed with 400 µL of 1 M perchloric acid. The deproteinized supernatant was neutralized by 3 M KHCO_3_. The ATP and acetyl-CoA assays were performed as described in the *in vitro* section.

### Electron microscopy studies

RAW 264.7 macrophages exposed to 2-AA and corresponding controls were fixed with 2% glutaraldehyde in 0.1 M cacodylate buffer and postfixed in 1% OsO4 in 0.1 M cacodylate buffer for 1 h on ice. The cells were stained all at once with 2% uranyl acetate for 1 h on ice, after which they were dehydrated in a graded series of ethanol (50–100%) while remaining on ice. Ultrathin (70 nm) sections were cut using a Leica EMUC7 ultramicrotome and collected onto formvar-coated grids (EMS, Hatfield, PA). Sections were contrast-stained using 2.0% aqueous uranyl acetate. Grids were examined at 80 kV in a JEOL 1011 transmission electron microscope (Peabody, MA) equipped with an AMT digital camera and proprietary image capture software (Advanced Microscopy Techniques, Danvers, MA).

## Author Contributions

Author contributions: L.G.R. conceptualization, A.B., A.C. and L.G.R., designed research; A.B., A.C. V.K.S., S. C., performed research; A.B., A.C. V.K.S., F.K., and L.G.R. analyzed data; W.M.O and A.A.T., provided resources, and F.K. and L.G.R. wrote the paper.

## Classification

Biological, health, and Medical Sciences/Microbiology

## Acknowledgments and Funding sources

We thank Dr. Vamsi Mootha and Sneha Rath for access and help with using the Seahorse instrument, respectively. We also thank Dr. Diane Capen and Dr. Dennis Brown for the guidance and processing of the ultramicroscopy images. This work was supported by the NIH award R01AI134857, The John Lawrence Massachusetts General Hospital Research Scholar Award and Shriner’s grant 83009 to L.G.R., and the Shriner’s grant 85132 to A.A.T. The funders had no role in the study design, data collection, analysis, decision to publish, or manuscript preparation.

---

**Figure S1.**
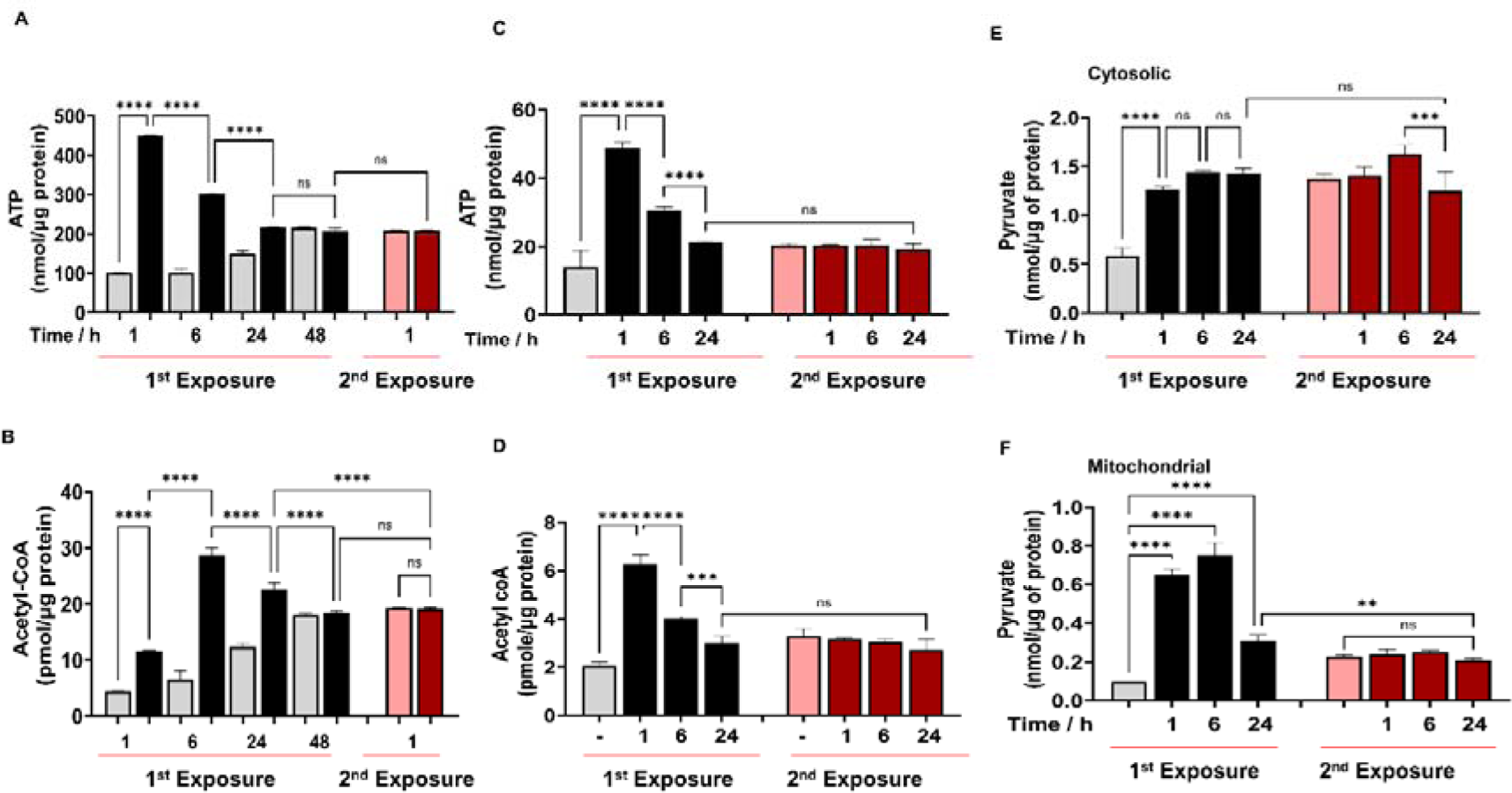
2-AA tolerization decreases metabolites in murine RAW 264.7 (A-B) and human THP-1 (C-D) cells. (A and C) ATP, (B and D) acetyl-CoA, and (E) cytosolic and (F) mitochondrial pyruvate levels were quantified in 2-AA (400 μM) exposed and re-exposed cells for the hours indicated using the same condition as shown in Fig. 1A. Mean ± SD is shown (*n* = 3); *** *p* < 0.001, and ns indicates no significant difference. One-way ANOVA followed by Tukey’s post hoc test was applied. The same color code, black cells after 1^st^ exposure and red cells after 2^nd^ exposure, was kept throughout the main manuscript, with corresponding controls shown in grey and pink, respectively.

**Figure S2:**
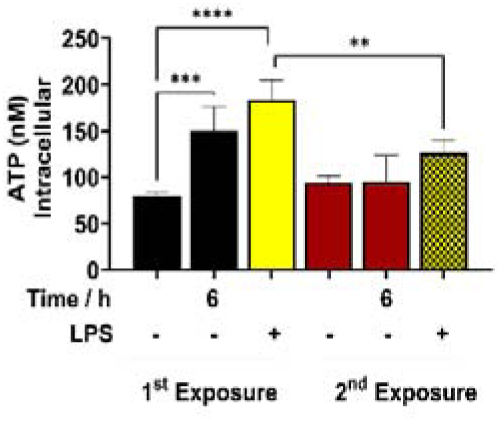
ATP levels in murine BMDM (A) cells with and without 2-AA (400 μM) or LPS stimulation (100 μg/mL). Each dot represents 1 experimental replicate of 4 independent experiments (*n* = 4). Data are presented as mean ± SD, \*\**p* < 0.01, \*\*\**p* < 0.001, and ns indicates no significant difference. One-way ANOVA followed by Tukey’s post hoc test was applied.

**Figure S3:**
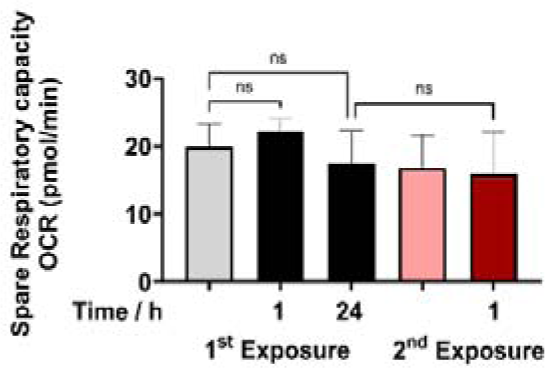
Effect of 2-AA on mitochondrial spare respiratory capacity in BMDM non-tolerized (black) and tolerized (red) macrophages. Controls are shown in grey and pink, respectively. Mitochondrial spare respiratory capacity data extrapolated from the OCR profiles shown in Figure 1D. Unstimulated BMDM macrophages were used as a control (c). Means ± SDs are shown, *n* = 6, * *p* < 0.05, *** *p* < 0.001, and ns indicates no significant difference. One-way ANOVA followed by Tukey’s post hoc test was applied.

**Figure S4:**
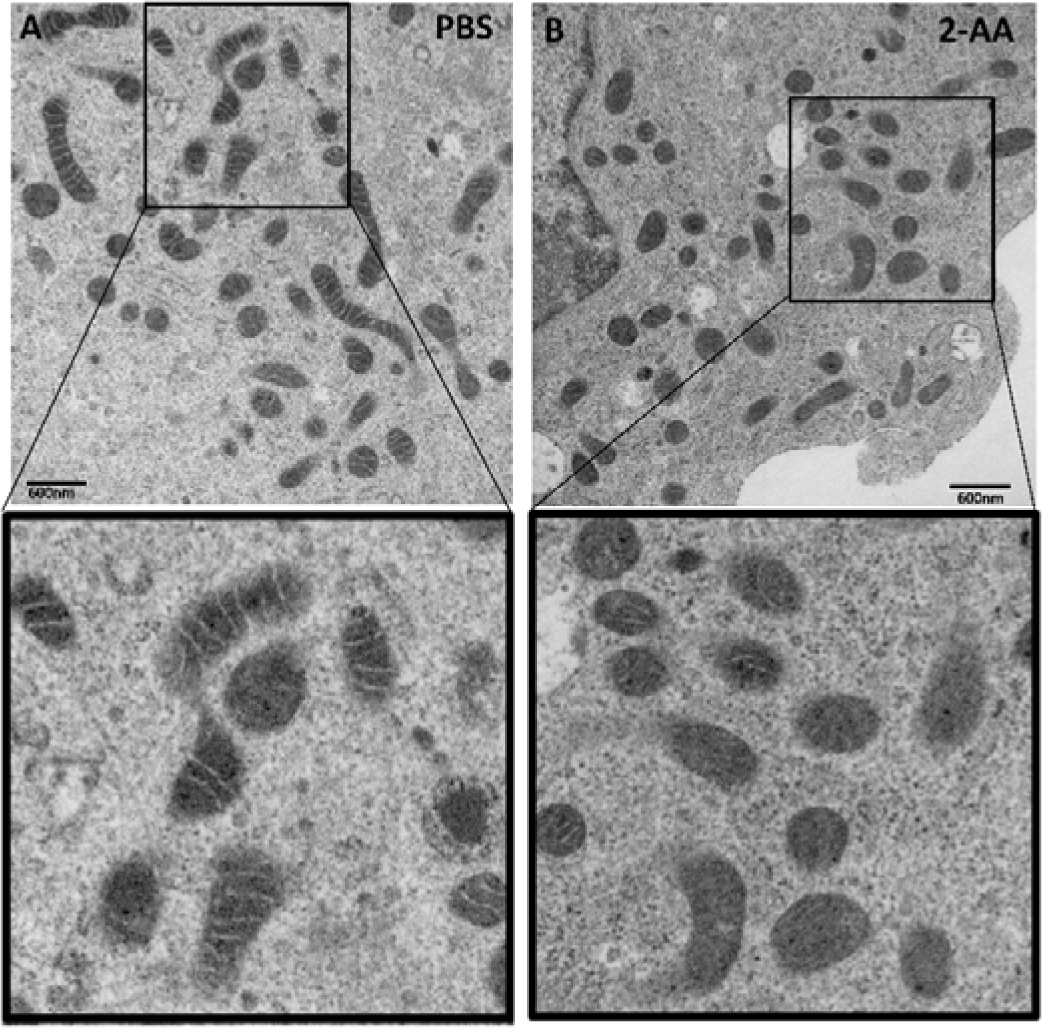
Transmission electron microscopy (TEM) images showing structural alterations of mitochondria in 2-AA exposed macrophages for 48 h. High-magnification (30,000X) TEM images showing mitochondrial abundance and structural changes in 2-AA exposed macrophages as compared to the non-exposed macrophages.

**Figure S5:**
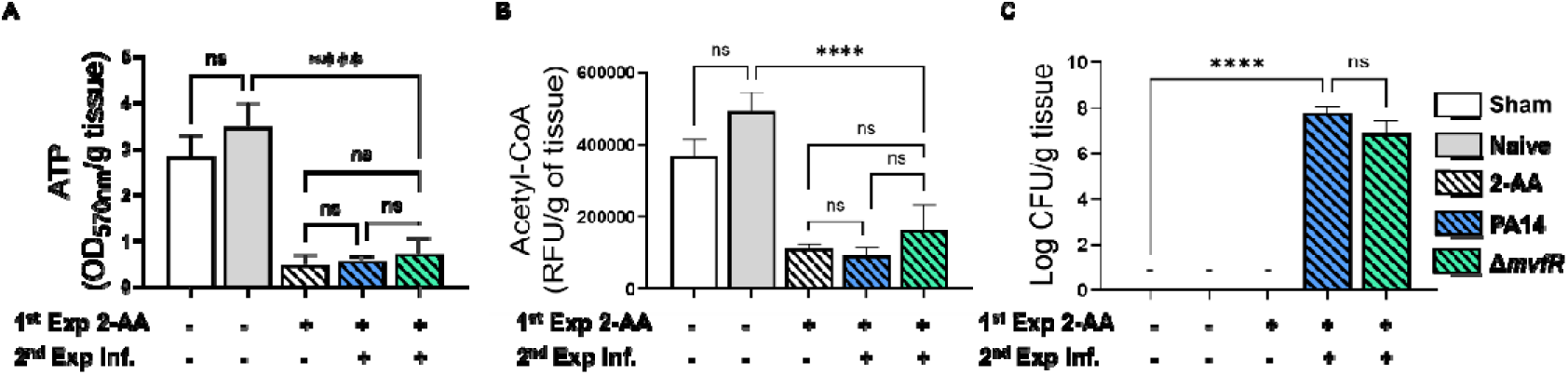
Exposure and re-exposure to 2-AA promotes a long-lasting decrease in ATP and acetyl-CoA levels and sustains bacterial presence in mice receiving 1^st^ exposure to 2-AA by injecting 2-AA and 2^nd^ exposure through infection with PA14 or Δ*mvfR* four days post-2-AA injection. (A) ATP and (B) acetyl-CoA concentrations in the spleens of mice. (C) Bacterial burden in muscles expressed as CFU counts was analyzed using the Kruskal–Wallis non-parametric test with Dunn’s post-test; \*\*\**p* < 0.001, and ns indicates no significant difference. Control mice groups: naïve were not given 2-AA; mice receiving 2-AA were given a single intraperitoneal injection of 2-AA four days prior to infection; sham represents a burn/PBS group since the burn and infection model was used. N = 4 mice per group, and 10 days post-infection is shown. Means ± SD are shown, \**p* < 0.05, \*\**p* < 0.01, \*\*\**p* < 0.001, and ns indicate no significant difference. One-way ANOVA followed by Tukey’s post hoc test was applied.

